# Modulating Subjective Pain Perception with Decoded MNI-space Neurofeedback

**DOI:** 10.1101/2023.10.25.563972

**Authors:** Taryn Berman, Cody Cushing, Shawn Manuel, Étienne Vachon-Presseau, Aurelio Cortese, Mitsuo Kawato, Choong-Wan Woo, Tor D. Wager, Hakwan Lau, Mathieu Roy, Vincent Taschereau-Dumouchel

## Abstract

Pain is a complex emotional experience that still remains challenging to manage. Previous functional magnetic resonance imaging (fMRI) studies have associated pain with distributed patterns of brain activity (i.e., brain decoders), but it is still unclear whether these observations reflect causal mechanisms. To address this question, we devised a new neurofeedback approach leveraging real-time decoding of fMRI data to test if modulating pain-related multivoxel fMRI patterns could lead to changes in subjective pain experience. We first showed that subjective pain ratings can indeed be accurately predicted using a real-time decoding approach based on the stimulus intensity independent pain signature (SIIPS) and the neurologic pain signature (NPS). Next, we trained participants in a double-blinded decoded fMRI neurofeedback experiment to up- or down-regulate the SIIPS. Our results indicate that participants can learn to down-regulate the expression of SIIPS independently from NPS expression. Importantly, the success of this neurofeedback training was associated with the perceived intensity of painful stimulation following the intervention. Taken together, these results indicate that closed-loop brain imaging can be efficiently conducted using *a priori* fMRI decoders of pain, potentially opening up a new range of applications for decoded neurofeedback, both for clinical and basic science purposes.

## Introduction

To this day, pain often remains a challenge to manage. The challenge comes in no small part from the way pain is represented in the human brain. Indeed, previous studies have shown that the neural representation of pain is complex and widely distributed [1,2]. This poses a serious challenge for intervention as few techniques can readily modulate such distributed patterns of brain activity. One approach that could potentially help us achieve this feat is decoded functional magnetic resonance imaging (fMRI) neurofeedback [3]. Decoded fMRI neurofeedback is a closed-loop brain imaging approach that combines machine learning with fMRI neurofeedback [3]. Several studies using this approach have shown that individuals can be trained to alter precise patterns of activity in their brain [3–8]. This specific type of neurofeedback is thought to operate following the basic principle of reinforcement learning, meaning that a rewarded “action” – modulating a pattern of neural activity – is in turn reinforced [9]. In this view, neurofeedback may not differ significantly from any type of skill learning. The main difference being that the rewarded action – changing patterns of neural activity – is concealed and requires “decoding” in order to be reinforced.

One of the main potential advantages of neurofeedback for therapy is that patients could learn to regulate their pain at will and without any medication. From the point of view of patients, they would simply learn to optimize reward by, unknowingly, regulating the real-time prediction of a pain decoder [10,11]. Previous experiments showed that such interventions can be conducted completely unconsciously and still present very specific effects on physiology and behavior [4–8,12]. For instance, it was shown that facial preferences can be modulated in the positive or negative direction by training participants to regulate brain activity within the cingulate cortex [12]. Following this rationale, it might be possible for participants to regulate their pain-related multivoxel patterns without providing any explicit regulation strategies. Demonstrating control over pain reports in this context would represent a rigorous double-blind test of the direct link between the multivoxel patterns and pain experience, expanding previous findings reported using univariate neurofeedback [13].

Before neurofeedback therapy can be easily and widely implemented, several objectives must be achieved. Firstly, we need a clear target for neurofeedback as the intervention’s success depends directly upon our capacity to correctly identify the relevant pattern of brain activity. Most previous decoded neurofeedback approaches aimed at training new brain decoders for each individual participant. This approach likely affords the most individualized interventions but also presents the disadvantage of being lengthy, as many experimental trials need to be acquired for each subject in order to build new individualized decoders. With reduced data, individualized brain targets tend to be more “noisy” and less accurate [14].

Between-subject decoders trained in the Montreal Neurological Institute (MNI) space on independent participants can potentially circumvent this problem [6]. It was previously shown that such between-subject decoders can sometimes outperform individualized ones [15,16]. Over the last decade, two pain “signatures” were developed that can accurately predict pain ratings in out-of-sample individuals [1]. The first one – the neurologic pain signature (NPS) – was trained to track pain triggered by nociceptive stimuli [2]. The second one – the stimulus intensity independent pain signature (SIIPS) – was developed specifically for predicting variations in pain ratings that cannot be explained directly by the nociceptive input [1] (see Figure 1A). This was achieved by training the decoder to predict the variations in residual pain ratings after controlling for the stimulation intensity and NPS response. For this reason, the SIIPS decoder is thought to capture higher-level psychological factors – such as placebo and cue-induced expectations [1] – which are presumed to contribute to the perception of pain. While the SIIPS and NPS decoders appear to be sensitive and specific to the pain experience [1,2], these findings remain correlational. As a result, it is still unclear if modulating the expression of these decoders could also change the associated subjective experience. Answering this question could potentially help narrow our search for new treatment strategies and help us directly target how patients subjectively feel [17].

**Figure 1.**
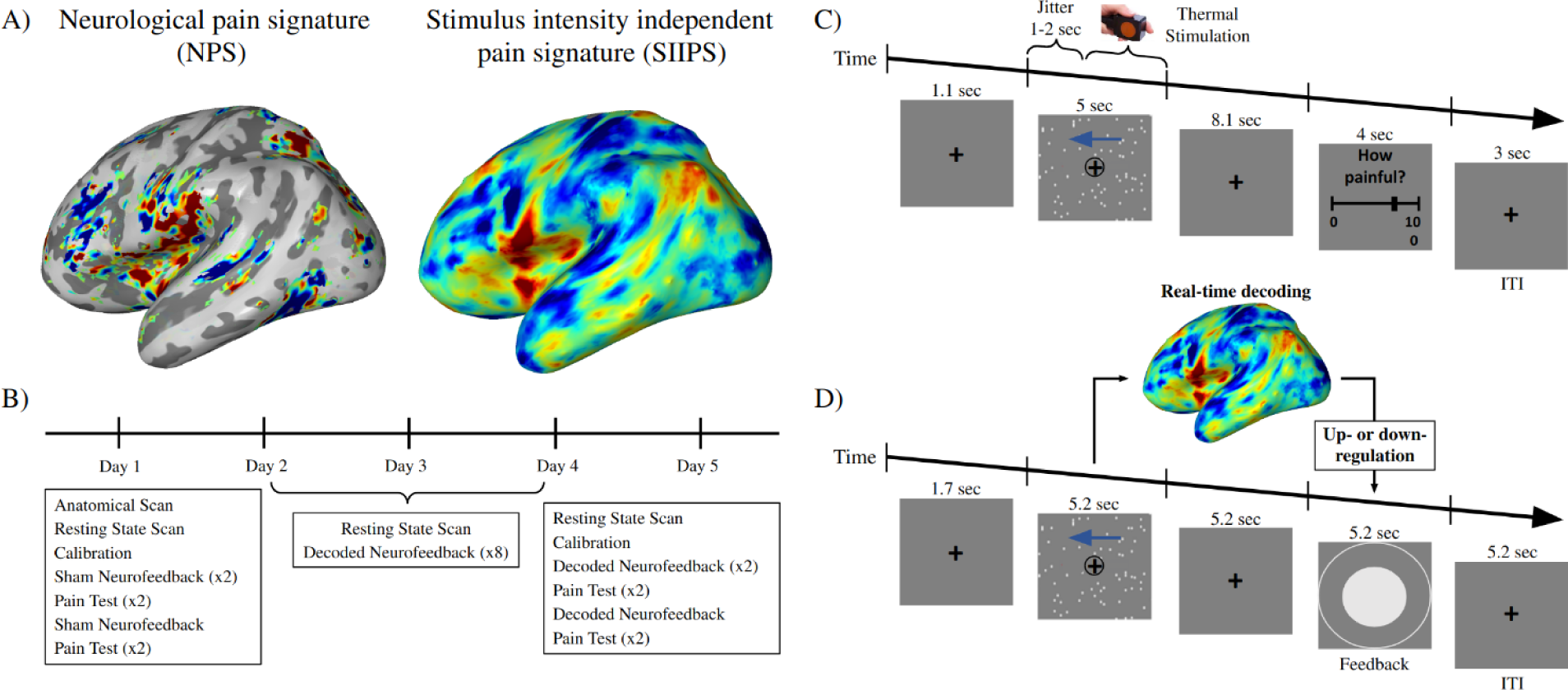
A) Model weights of the Neurological pain signature (NPS) and Stimulus independent pain signature (SIIPS). B) Study outline indicating the experimental procedure. C) One trial of the pain assessment sessions (“pain session”) conducted before and after the decoded neurofeedback intervention. D) One trial of decoded fMRI neurofeedback.

However, using an “off-the-shelf” decoder trained in the MNI space poses some technical challenges. This requires a different approach from previous decoded neurofeedback experiments, which were predominantly conducted in the native space using decoded brain activity from only a few hundred voxels. Here, our goal is to process and transform whole-brain data into the MNI space in real-time in order to conduct decoding with ∼200,000 voxels. To address this challenge, we adapted the ATR toolbox (https://bicr.atr.jp/decnefpro/software/) to create a new real-time analytical pipeline by implementing most computations in parallel using the graphic processing unit (GPU) of a relatively standard desktop computer. We first determined if this processing pipeline could accurately predict pain ratings in real-time in out-of-sample participants using SIIPS and NPS. Then, we randomly assigned participants to be rewarded for the up- or downregulation of SIIPS. We conducted a five-day double-blinded decoded neurofeedback experiment to assess the effect of training on subsequent pain ratings. First, we hypothesized that participants in the up- and downregulation groups will respectively increase and decrease the expression of SIIPS during neurofeedback, without changing the expression of NPS, a brain decoder reflecting a related but distinct aspect of pain perception. Second, we hypothesized that the modulation of pain ratings will be proportional to the success of neurofeedback training.

To anticipate, we show that we can accurately predict pain ratings in real-time in out-of-sample participants using SIIPS and NPS. Furthermore, we show that participants can be trained to down-regulate the expression of SIIPS and that the success of this training is associated with the intensity of perceived pain following the neurofeedback training.

## Methods

### Participants

Twenty-two participants were recruited from the McGill Psychology Human Participant Pool, the McGill Studies for Cash Facebook page, and the Concordia Studies for Cash Facebook page. Of those recruited, three withdrew immediately after signing up and three were excluded because they did not meet the inclusion criteria. The final sample included 16 participants who ranged in age from 19-29 years (*M* = 21.62; *SD* = 2.94). The study was conducted at the Montreal General Hospital magnetic resonance imaging platform. Inclusion criteria were: (a) aged between 18 and 45; (b) no current diagnosis of a neurological, psychiatric, or pain-related disorder; (c) no psychotropic medications; (d) no contraindication to magnetic resonance imaging. These inclusion criteria were specified on the recruitment advertisements and verified through screening forms and an additional assessment on the first day of the study. The final sample contained nine (56.25%) women and seven (43.75%) men. The study was approved by the McGill University Research Ethics Board and the participants provided informed consent.

### MRI parameters

Participants were scanned using a 3T MRI scanner (i.e., Prisma Siemens) at the Montreal General Hospital with a 64-channel head coil. During the experiment, we obtained 33 contiguous slices (TR = 0.867 s, TE = 20 ms, Acceleration factor PE = 2, voxel size = 3 × 3 × 3 mm ^3^, field-of-view = 192 × 192 mm, matrix size = 64 × 64, slice thickness = 3 mm, 0 mm slice gap, flip angle = 58 deg) oriented parallel to the AC-PC plane, which covered the entire brain. We also obtained T1-weighted MR images (MP-RAGE; 256 slices, TR = 2250 ms, TE = 3.06 ms, 4 voxel size = 1 × 1 × 1 mm ^3^, field-of-view= 256 × 256 mm, matrix size = 256 × 256, slice thickness = 1 mm, 0 mm slice gap, TI = 900 ms, flip angle = 9 deg).

### Study Design

Participants were blinded to experimental conditions and randomly assigned to either up-(*N* = 7) or downregulate (*N* = 9) the pain decoder (i.e., SIIPS [1]). The study outline is shown in Figure 1B. The experiment was conducted across five days, with days two to five occurring consecutively. A resting state scan was conducted at the beginning of each day to determine the spontaneous fluctuations of SIIPS at rest. This information was used to establish a baseline to scale feedback in real time using the mean and standard deviation of the distribution (see the *Real-time Processing* section). Day one included an anatomical scan, a pain calibration session, and the first pain session. We interspersed two blocks of sham neurofeedback between the pain session blocks in order to match the sequence of events in the post-test procedure (Figure 1B). Days two to four consisted of decoded neurofeedback training based on SIIPS activity. On day five, participants performed a calibration session, three neurofeedback blocks (interspersed between the pain session blocks), and the second pain session to determine whether the intervention changed their subjective pain ratings.

#### Calibration Session

A calibration session was conducted in the fMRI scanner before each pain session (i.e., on days one and five) and determined the stimulus intensity that corresponded to a pain rating of 50/100 on the visual analog scale. This was achieved by delivering 14 stimulations of seven intensities (i.e., 41, 45, 47, 48, 48.5, 49, 49.5 degrees celsius), each lasting 3 seconds, with a ramp up speed of 75 degrees/second (administered using the five bars of an MR compatible T 09 probe and TCS device of QST.lab). We collected subjective pain ratings following these stimulations and fitted an exponential function to the data in order to infer the stimulation intensity associated with a pain rating of 50 on a visual analog scale ranging from zero (i.e., no pain) to 100 (i.e., worst pain imaginable).

#### Pain Session

The pain sessions consisted of applying painful thermal stimulation to the left forearm during the presentation of the target visual cue (i.e., leftward dot motion) or control visual cue (i.e., still image of a fixation cross surrounded by a circle; see Figure 1D). Participants were asked to report their pain rating after each trial using a visual analog scale, which ranged from zero (i.e., no pain) to 100 (i.e., worst pain imaginable). They performed four blocks, with 12 trials per block. Six of the trials included leftward dot motion (i.e., target visual cue) while the other six included the still image (i.e., control visual cue). These conditions were included in order to test if an association could be established between the visual cue and the SIIPS pattern (see Condition in Table 3). The intensity of painful stimulations remained constant across all pain trials and corresponded to the stimulation intensity established during the calibration session.

#### Decoded Neurofeedback Training

In each trial of decoded neurofeedback training, the target visual cue (i.e., dots moving leftward) was presented during the induction period while the brain activity was being recorded (see Figure 1C). This data was preprocessed online (see *Real-time Processing* section) and SIIPS was applied to the averaged brain activity during the induction period in order to give a predicted pain value that was then presented visually to participants.

This feedback showed participants their accuracy in achieving the desired brain state based on the diameter of the inner circle. To make the task more engaging, whenever the brain activity was in the top 10 percentiles of expected expression (see *Real-time Processing* section), the feedback circle turned blue indicating a “high score” bonus. Similarly, when scores were in the top 30 percentiles for 3 trials in a row, a “high streak” bonus was provided. Importantly, participants were not provided with any explicit training strategy; they were only asked to manipulate their brain activity in order to maximize their monetary reward [10,11]. The experiment was conducted in a double-blind setting; neither the participants nor the experimenters were informed of the exact nature of the target decoder. They were simply informed that they could receive up to an extra 20 dollars after each training day if they performed well on the task. Participants performed eight blocks of 16 trials on each of the three training days (i.e., days two to four), for a total of 128 trials per day.

### Real-time Processing

It is necessary to conduct real-time analyses quickly in order to provide participants with feedback regarding their brain activity. The feedback was presented six seconds after the end of the induction cue in order to account for the delay of the hemodynamic response function. An innovative aspect of the current study was to develop an analytical pipeline that included computations carried with graphical processing units (GPU). Typical analytical pipelines conducted on the CPU can be quite lengthy when applied to whole-brain data. For this reason, we devised a GPU implementation that included the realignment procedure of the BROCCOLI toolbox [18] as well as data management, standardization, detrending, and baseline correction (see Figure 2-3) conducted using the MATLAB GPU array. The normalization and smoothing of the data was carried out using functions from Statistical Parametric Mapping (SPM) 12 (www.fil.ion.ucl.ac.uk/spm) [19].

**Figure 2.**
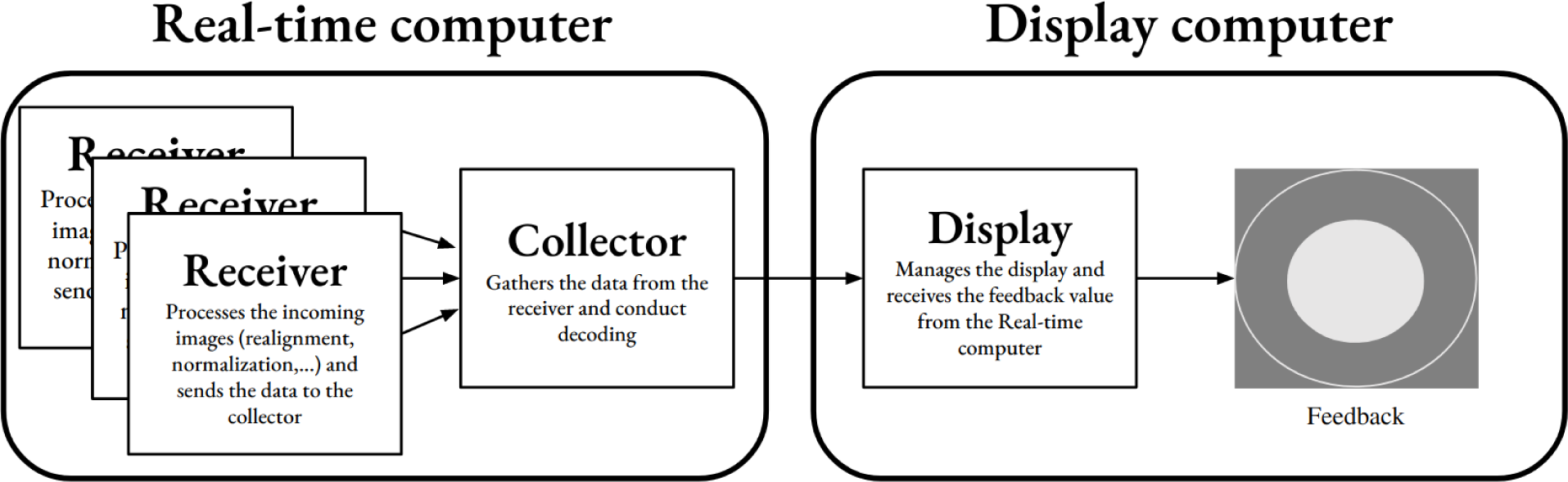
Schematic representation of the real-time processing procedure. Brain data are first transferred from the MRI console to the real-time processing computer through a local network. The “receiver” instances pre-process each incoming image and send the processed data to the “collector” instance which, in turn, is responsible for detrending and decoding the data (see main text for more detail). The result of the decoding procedure is then communicated through the local network to the display computer which is responsible for managing the experiment display and providing the visual feedback to participants.

**Figure 3.**
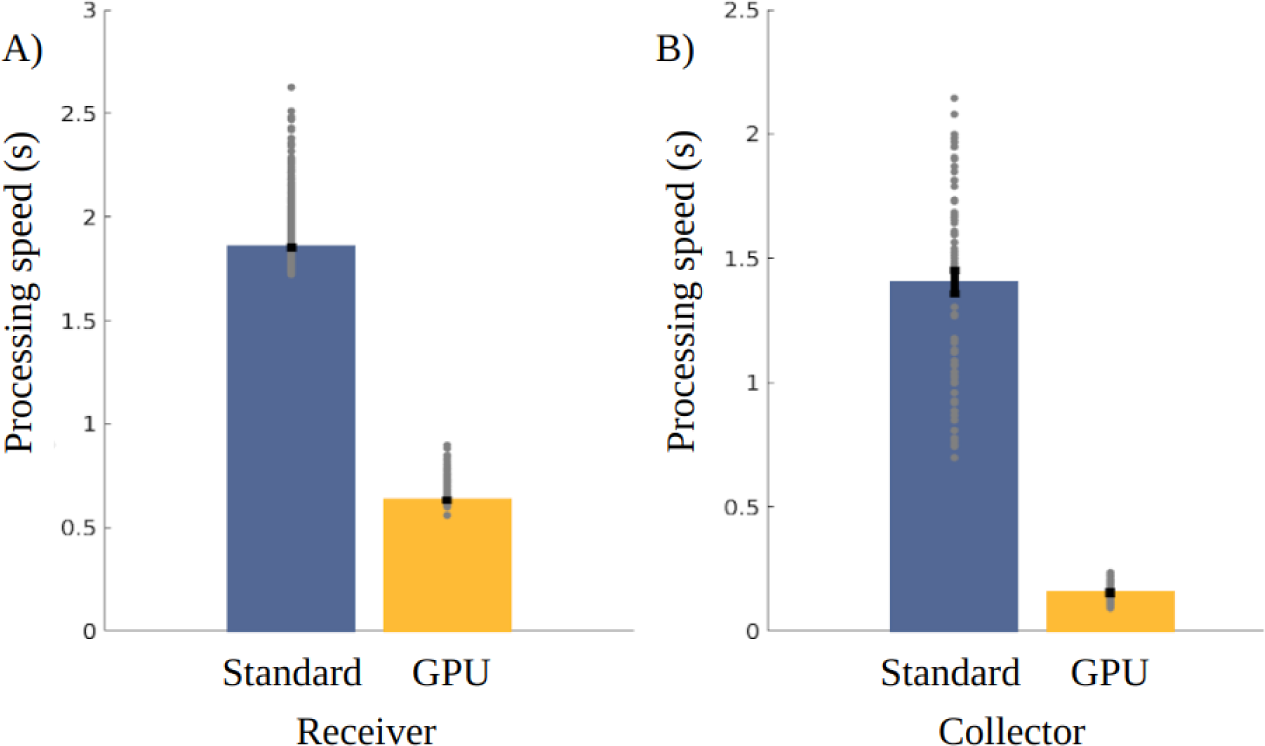
Speed of real-time processing of fMRI data by (A) the “receiver” and (B) the “collector” instances. Each data-point corresponds to one MRI image. Overall, the GPU implementation allowed processing and decoding data in 0.78 second compared to 3.25 seconds using the standard approach. Error bars represent standard error of the mean.

There were three “receiver” instances that processed DICOM images as soon as they were written to the real-time processing computer. These receivers converted the DICOM images to a NIfTI format, realigned the NIfTI image to a mean fMRI BOLD image, normalized the images to the MNI space and then sent this data to another MATLAB instance which can be called the “collector”. This instance gathered pre-processed data from all three receivers. The collector further processed the data by removing linear trends from the fMRI activity (using a 20 second baseline at the beginning of the experiment) and *z*-scoring it voxel-wise. To account for the hemodynamic response function, we shifted the data by 6 TRs (TR = 0.867 s), averaged the signal within a 6-TR time window and submitted this pattern to SIIPS. Since the output of SIIPS cannot be directly interpreted as reflecting a specific pain rating, we needed to scale the output in order to provide meaningful feedback to participants. In order to do so, we ran SIIPS on resting state data to obtain a distribution of SIIPS expression at rest. We then used this distribution to scale the feedback. Specifically, we obtained a percentile for the current SIIPS expression based on the mean and standard deviation of the distribution that was observed during the resting state session on the same day. In order to make the feedback more explicit to participants, percentiles were rescaled so that values below 30 and above 70 were rescaled to 0 and 100 respectively (Feedback score = (current percentile - 30)/(70-30)). This feedback score was communicated to participants through the diameter of the feedback circle and paired with monetary reward.

Participants were informed that the diameter of the circle was directly proportional to their monetary reward and that they could make up to an extra 20$ per day. In the up-regulation group, the feedback score was directly proportional to the diameter of the circle while in the downregulation group, the feedback score was inverted (100 - feedback score). This was achieved so that a decreased SIIPS expression would be paired with the monetary reward. The scalar value obtained using this procedure was transferred through the local network to another MATLAB instance responsible for controlling the display presented to participants.

## Data Analysis

### Real-time processing speed and pain decoding

We compared the processing speed of the real-time approach implemented on the GPU to a standard approach using exclusively the central processing units (CPU). We compared the processing speed of the “receiver” (i.e., alignment and normalization) and “collector” instances (i.e., detrending, standardization and decoding). This was achieved using paired sample *t*-tests comparing the processing speed of individual DICOM images and decoding trials.

We tested the real-time decoding procedure by evaluating the capacity of SIIPS and NPS to predict pain ratings during the calibration session. As described in the study design, we conducted a calibration session where participants received painful stimulations of varying intensities and were asked to provide pain ratings after each stimulation. We used SIIPS and NPS to obtain predicted pain scores based on brain activity and correlated the predicted scores with the subjective ratings of participants. We conducted a sliding window analysis (6-TRs) on the preprocessed images in order to determine how accurate the pain decoders were to predict subjective pain ratings. For each time point, we performed a within-subject correlation between the predicted and real pain ratings. One-sample *t*-tests were used to determine if the Fisher-transformed correlation coefficients were different from zero. Significance was established using a permutation test (i.e., 5000 sign permutations over the tested time windows using a two-tail approach).

### Induction success

We used multilevel modeling to examine if each group was able to successfully modulate SIIPS induction. Furthermore, we evaluated if NPS activity (which was not the target of training) was also affected by the intervention. In this model, trial and log(trial) are included as level 1 predictors in order to control for habituation/sensitization, while day (1 to 4) is at level 1 to control for changes in induction success across time. In this model, group (up- or downregulation) and decoder type (SIIPS or NPS) are level 2. Trial and log(trial) are grand mean centered, group and decoder type are effect coded, and day is continuous. Bootstrapped estimates were computed using the *tab_model* function in the *sjPlot* package [20] for all models based on bootstrapped distributions (10000 replications, resampled at the trial level). The same function was used to compute marginal and conditional R^2^ values. All multilevel models were built using fixed predictors that should influence the outcome variable based on theory. Random effects were added if they improved model fit.

The multilevel model can be represented by the following equations:

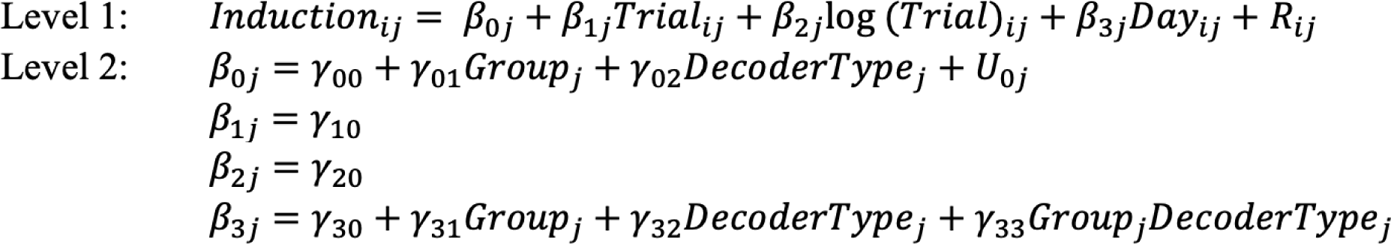

In the above equations, induction success (*Induction_ij_*) for each participant (denoted by *j*) on each trial (denoted by *i*) is predicted by the level 1 variables trial (γ_10_), log(trial) (γ_20_), and day (γ_30_), and the level 2 variables group (γ_01_) and decoder type (γ_02_). There are also cross-level interactions examining groups across days (γ_31_), decoder type across days (γ_32_), as well as group and decoder type across days (γ_33_). This model also contains the fixed component of the intercept (γ_00_), a random component which is the intercept variance for each person (*U_0j_*) with total variance *τ*^2^, and the level 1 residual (*R_ij_*) with variance σ^2^.

In order to further investigate the interactions, we next ran two different multilevel models - one for each decoder - to determine if participants were able to modulate SIIPS and NPS induction separately across days. In these models, trial, log(trial), and day are level 1 predictors, while group is level 2. The following equations represent the models:

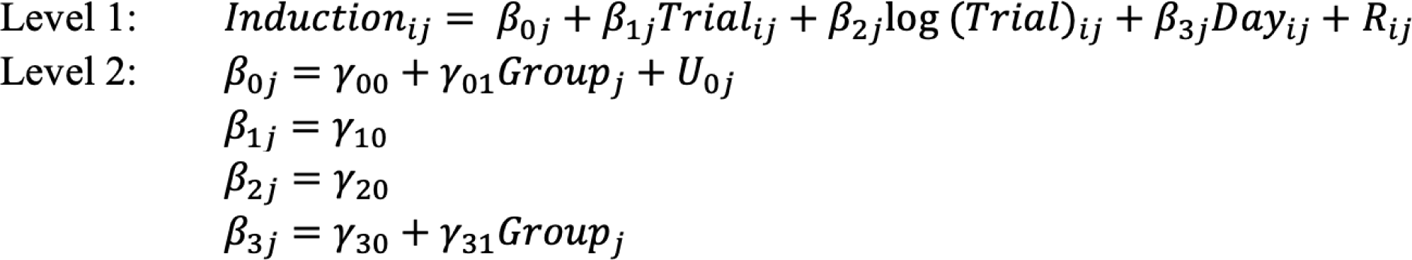

In the above equations, SIIPS and NPS induction (*Induction_ij_*) for each participant on each trial is predicted by the level 1 variables trial (γ_10_) log(trial) (γ_20_), and day (γ_30_), and the level 2 variable group (γ_01_). There is also a cross-level group by day interaction (γ_31_). These models also contain a fixed intercept (γ_00_), random intercept variance for each person (*U_0j_*) with total variance *τ*^2^, and a level 1 residual error (*R_ij_*) with variance σ^2^.

Next, we wanted to further examine if each group was able to separately modulate SIIPS induction. To examine this, two multilevel models were run: one for the upregulation and one for the downregulation group. These models can be represented by the below equations:

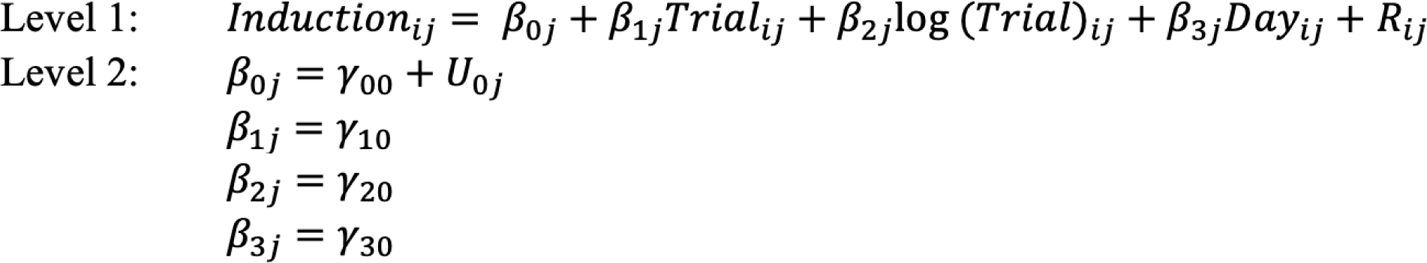

Here, SIIPS induction for each individual in each group is being predicted. All other variables are the same as in the previous models.

### Pain ratings

To ensure that no difference in pain ratings was present between groups on day one, an independent-samples *t*-test was run between groups on averaged pain ratings for each person on day one and 95% confidence intervals were computed. To determine if pain ratings on day five were related to SIIPS induction, we employed multilevel modeling. In this model, condition (i.e., whether or not dot-motion was viewed during induction), trial, and log(trial) were all level 1 predictors, while group, pre-rating (i.e., pain ratings from day one), and SIIPS induction were all level 2. The following equations represents this model:

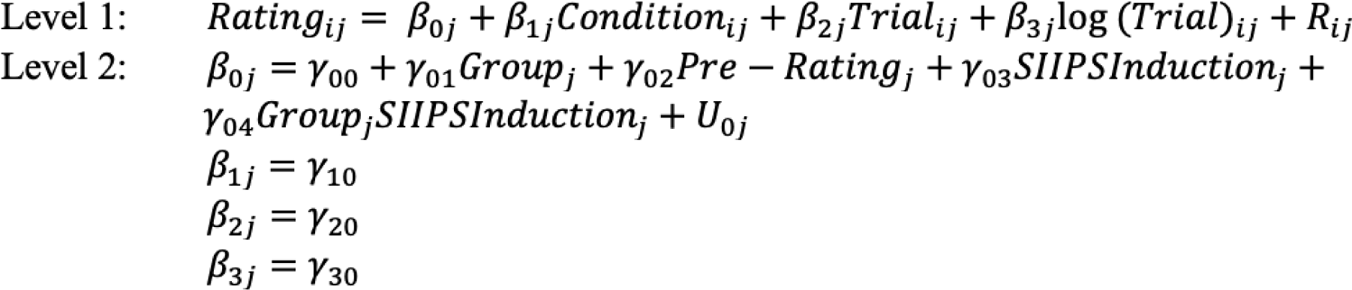

In the above equations pain ratings on day five (for each trial of each individual in each group) is being predicted by the level 1 variables condition (γ_10_), trial (γ_20_), and log(trial) (γ_30_), and the level 2 variables group (γ_01_), pre-rating (γ_02_), SIIPS induction (γ_03_), and group-by-SIIPS induction (γ_04_). This model also contains a fixed intercept (γ_00_), random intercept variance (*U_0j_*) with total variance *τ*^2^, and a level 1 residual error (*R_ij_*) with variance σ^2^.

To further investigate the interaction and group effect, two more models - one for each group - were run to determine if, in each group, SIIPS induction modulates pain ratings. These models can be represented by the following equations:

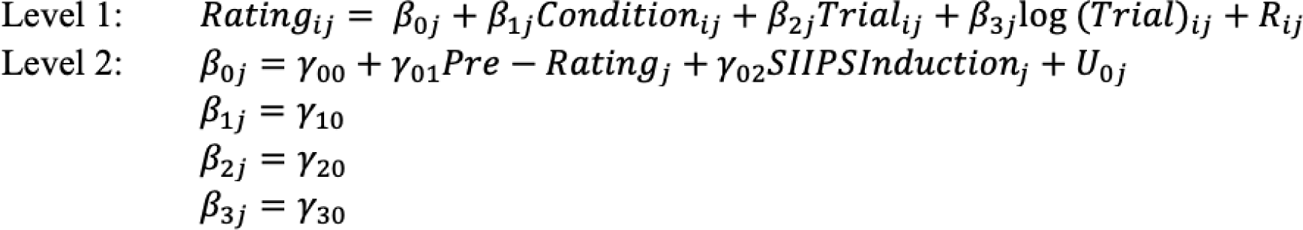

In the above equations, pain ratings on day five for each trial for each individual is being predicted. All other variables are the same as in the previous models. All multilevel analyses were conducted in the RStudio software for statistical analysis (RStudio Team, 2020, version 1.3.1056 [21] using the *lme4* package, version 1.1.32 [22]). Data and analysis scripts are available online (https://osf.io/mwvt5/?view_only=6633772dab064049887f6f9fe5b9188f).

## Results

### Real-time processing speed and pain decoding

The “receiver” processing speed was significantly faster using our GPU implementation (*M* = .63, *SD* = .03) than using the standard CPU approach (*M* = 1.85, *SD* = .09; *t*(2600) = -662.54, *p* < .001; Figure 3A). Similarly, the “collector” (i.e., detrending, standardization and decoding; Figure 3B) processing speed was also significantly faster using the GPU implementation (*M* = .15, *SD* = .04) than using the standard CPU approach (*M* = 1.4, *SD* = .4; *t*(69) = -28.78, *p* < .001). Overall, the GPU implementation allowed the processing of an fMRI image and decoding in 0.78 second compared to 3.25 seconds using the standard CPU approach.

We found that the prediction of SIIPS and NPS are indeed correlated with the pain ratings of participants (Figure 4). The correlation was the highest during the time window 6 TRs following the stimulation (SIIPS: *t*(15) = 3.843, *p* = .002, Cohen’s *d* = .99; NPS: *t*(15) = 9.049, *p* < .001, Cohen’s *d* = 2.33). These results bring validity to our real-time decoding procedure and justify using this time window as a target for neurofeedback.

**Figure 4.**
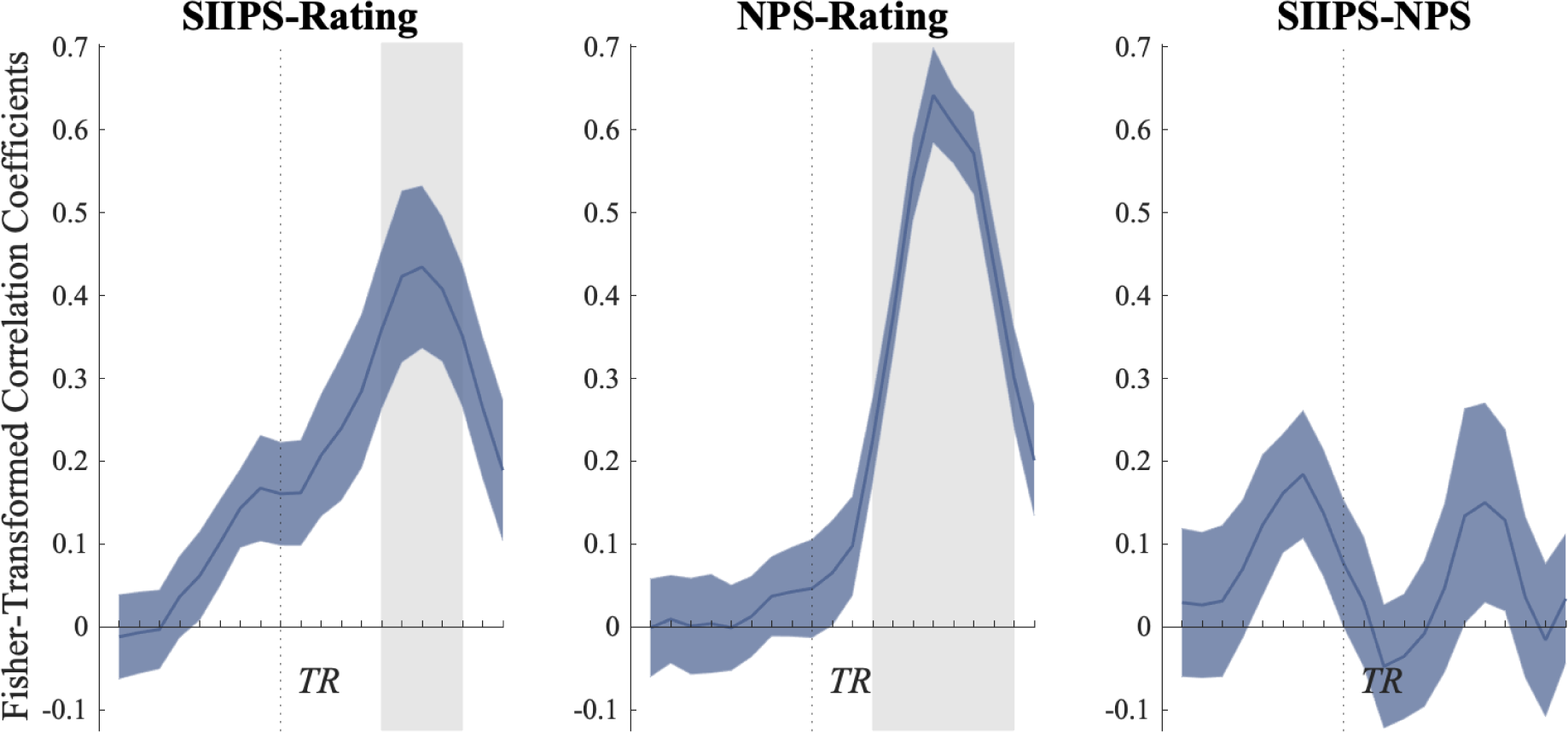
Correlations between the pain ratings and the real-time prediction of SIIPS (Left) and NPS (middle). As expected, the SIIPS and NPS decoders appear to explain independent parts of the variance of the subjective ratings (see right panel). Data were averaged within a 4-TR sliding window. The dashed lines correspond to the start of the thermal stimulation. For real-time decoding, we shifted the decoding window by 6 TR to account for the hemodynamic response function, which corresponds to the peak in decoding performance. Grey shaded area represents significant results based on a permutation test. Blue shaded regions indicate standard error.

### Induction Success

We ran a multilevel model to determine whether group membership had an effect on induction success (i.e., if participants are able to successfully modulate the activity of the SIIPS decoder independently from NPS activity, which was not the target of training). We hypothesized that participants could be trained to increase or decrease the expression of the SIIPS decoder as a function of the experimental group. Furthermore, we hypothesized that this modulation would be largely independent of the activity of NPS. We found a three-way interaction between decoder type (SIIPS or NPS), group (up- or downregulation) and day (Day 1 to 4; γ = 358.141 [66.942, 650.002], *t*(13256.00) = 2.375, *p* = .016; see Supplementary Material Table S1). Using separate models for SIIPS and NPS, our results indicate that group and day interact to predict SIIPSinduction (γ = 702.867 [121.826, 1283.2623], *t*(6621.25) = 2.339, *p* = .018; Table 1; Figure 5A), but not NPS induction (γ = -.010 [-.452, .431], *t*(6620.03) = -.053, *p* = .965; Table 2; Figure 5B). Specifically, it seems that participants in the downregulation group learned to decrease SIIPS expression over the course of training (γ = -632.032 [-868.812, -385.656], *t*(3732.06) = -5.137, *p* < .001; see Supplementary Material Table S2), while the participants in the upregulation group did not (γ = 875.656 [-442.724, 2194.109], *t*(2886.76) = 1.292, *p* = .199; see Supplementary Material Table S3). Therefore, these results demonstrate that the experimental manipulation was successful in teaching participants in the downregulation group to decrease the expression of SIIPS.

**Figure 5.**
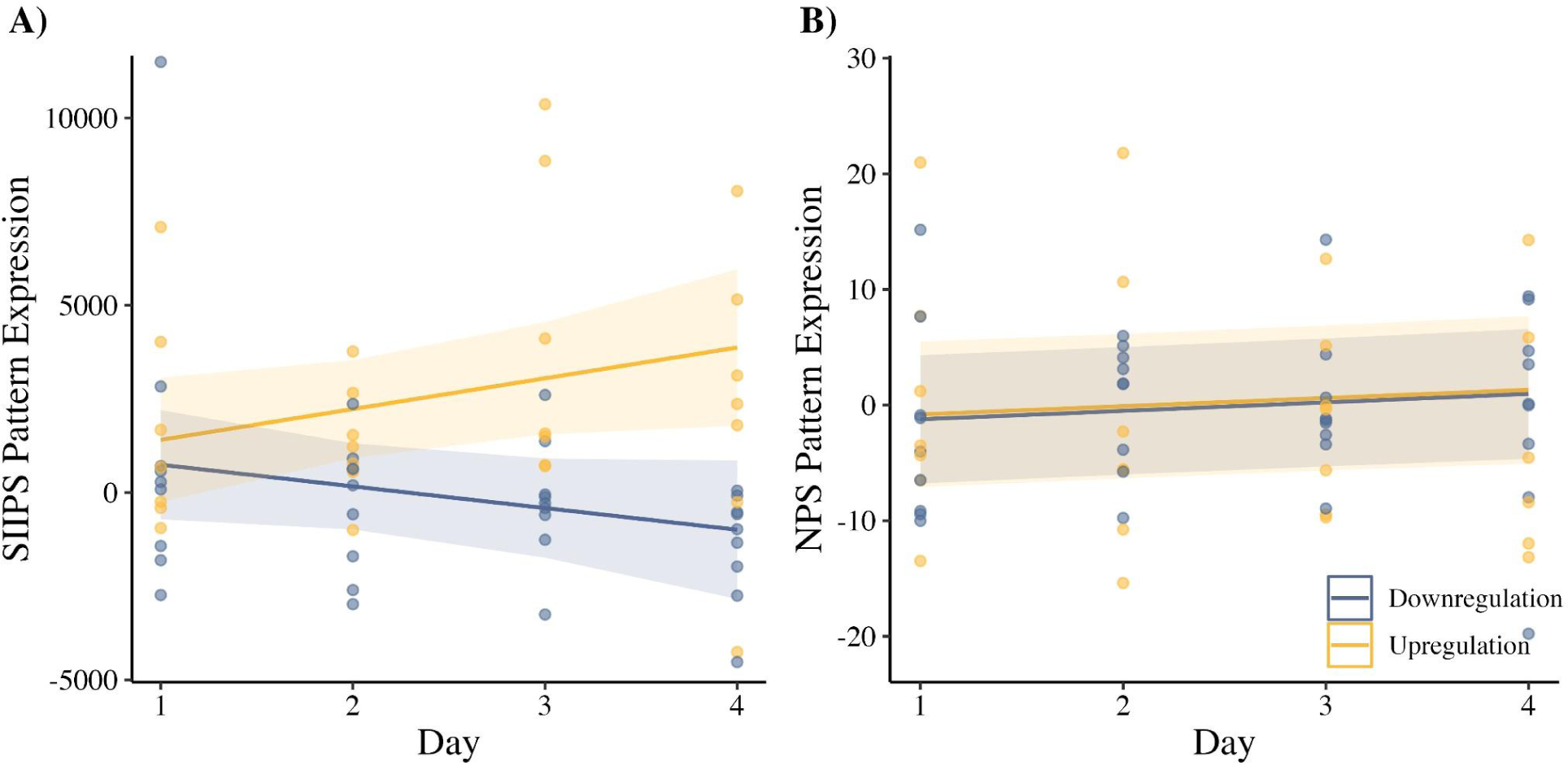
Multilevel model estimates of SIIPS (A) and NPS (B) decoder predictions across days for the upregulation (yellow) and downregulation (blue) groups. Data points indicate the mean decoder induction on each day for each individual participant. Shaded regions indicate standard error.

**Table 1.** *Multilevel model estimates for SIIPS induction. Estimates are bootstrapped 10 000 times.*

| *Predictors* | *Model Estimates* | *95% CI* | *p-values* | *Bootstrapped Estimates* | *Bootstrapped 95% CI* | *p-values* |
| --- | --- | --- | --- | --- | --- | --- |
| (Intercept) | 1076.631 | [-31.315, 2184.578] | .064 | 1091.116 | [-13.313, 2199.356] | .054 |
| Trial | 57.254 | [25.023, 89.485] | <.001 | 57.616 | [24.581, 89.367] | <.001 |
| log(Trial) | -1719.498 | [-2948.612, -490.384] | .006 | -1723.805 | [-2958.243, -483.144] | .006 |
| Group | 331.689 | [-767.691, 1431.069] | .554 | 325.133 | [-702.787, 1448.799] | .539 |
| Day | 121.410 | [-477.258, 720.078] | .691 | 119.332 | [-462.271, 715.766] | .687 |
| Group x Day | 699.958 | [113.339, 1286.576] | .019 | 702.867 | [121.826, 1283.263] | .018 |
| *Random Effects* |  |  |  |  |  |  |
| $\sigma^2$ | 524381716.22 | | | | | |
| $\tau_0^2$ | 1805223.43 | | | | | |
| Marginal/<br>Conditional<br>$R^2$ | .005/ .009 | | | | | |

*Note.* Group is effect coded where upregulation is 1 and downregulation is -1.

**Table 2.**
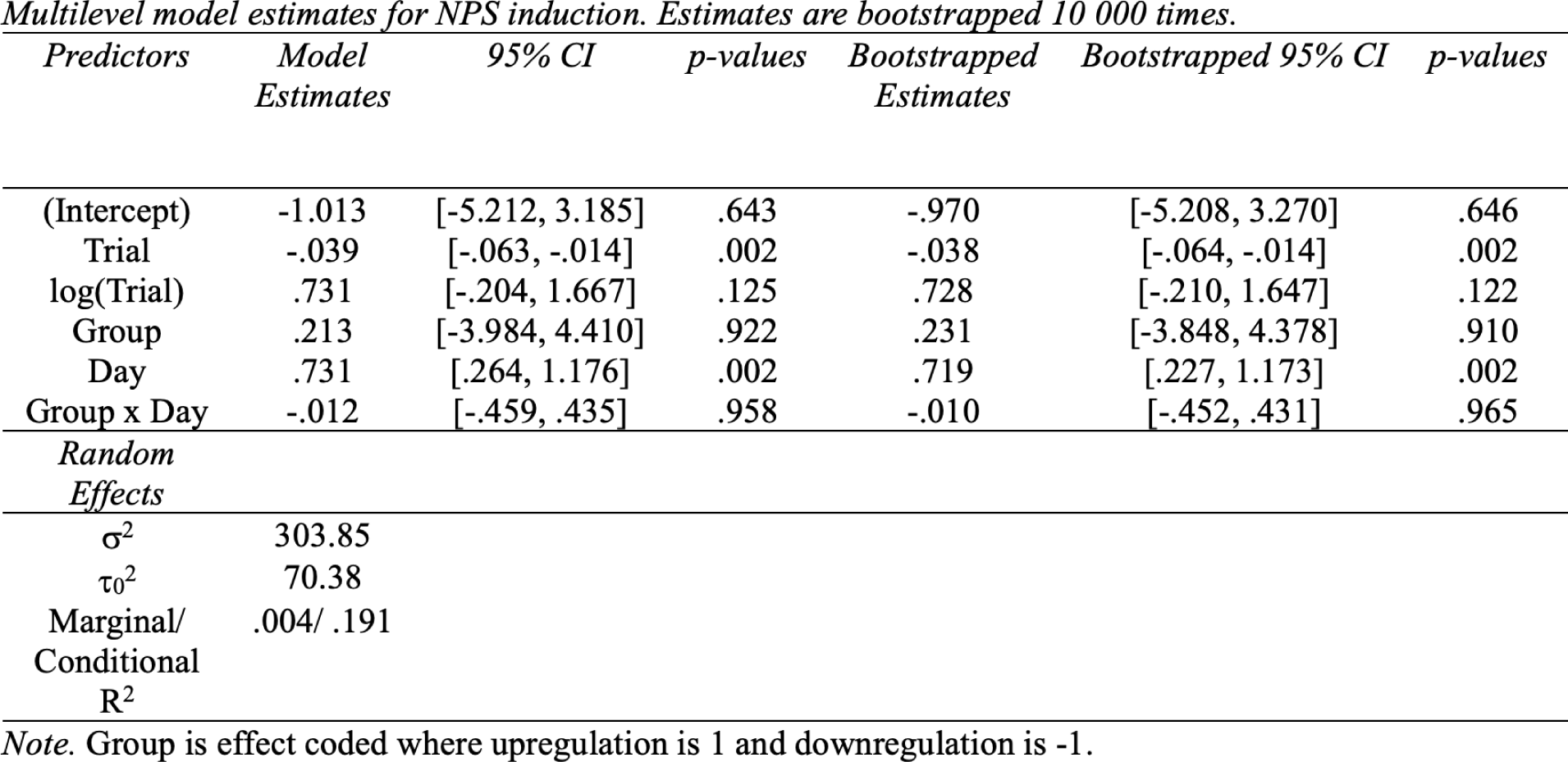

### Pain ratings

We investigated how the intervention affected the pain ratings of participants after neurofeedback training. There was no difference in pre-ratings between groups on day one (*M_up_* = 44.556 [36.789, 52.323], *M_down_* = 47.557 [41.426, 53.687], *t*(12.575) = -.5792, *p* = .573). We reasoned that group membership may impact pain ratings on day five as a function of neurofeedback success (see Table 3). Indeed, the results indicate a significant interaction between group and SIIPS pattern induction (γ = -13.981 [-20.878, -6.948], *t*(11.00) = -3.974, *p* < .001), such that in the downregulation group, a greater decrease in SIIPS induction was associated with a greater decrease in pain ratings (see Figure 6). When examining each group separately, we found that pain ratings were predicted by SIIPS induction in the downregulation group (γ = .009 [.004, .014], *t*(6.00) = 3.273, *p* = .001; see Supplementary Material Table S4), where less SIIPS induction was associated with lower pain ratings. In the upregulation group, SIIPS induction was not found to predict pain ratings (γ = .000 [-.002, .003], *t*(4.00) = .031, *p* = .963; see Supplementary Material Table S5).

**Table 3.** *Multilevel model estimates for pain ratings on day 5. Estimates are bootstrapped 10 000 times.*

| *Predictors* | *Model Estimates* | *95% CI* | *p-values* | *Bootstrapped Estimates* | *Bootstrapped 95% CI* | *p-value (based on bootstrapped estimates)* |
| --- | --- | --- | --- | --- | --- | --- |
| (Intercept) | 53.055 | [47.277, 58.833] | <.001 | 53.121 | [47.308, 58.908] | <.001 |
| Trial | 3.160 | [2.281, 4.038] | <.001 | 3.168 | [2.287, 4.039] | <.001 |
| log(Trial) | -19.596 | [-23.786, -15.405] | <.001 | -19.646 | [-23.801, -15.476] | <.001 |
| Condition | .611 | [-.371, 1.593] | .222 | .616 | [-.361, 1.576] | .212 |
| Pre-Rating | -.081 | [-.716, .554] | .807 | -.075 | [-.701, .554] | .813 |
| Group | -7.607 | [-14.467, -.746] | .052 | -7.525 | [-14.264, -.883] | .027 |
| SIIPS Induction | 14.155 | [5.590, 22.720] | .008 | 14.119 | [5.604, 22.591] | .001 |
| Group x SIIPS Induction | -13.932 | [-20.815, -7.049] | .002 | -13.981 | [-20.878, -6.948] | <.001 |
| *Random Effects* |  |  |  |  |  |  |
| $\sigma^2$ | 192.62 | | | | | |
| $\tau_0^2$ | 85.13 | | | | | |
| Marginal R<sup>2</sup>/<br>Conditional R<sup>2</sup> | .335/.539 |  |  |  |  |  |

*Note.* Group is effect coded where upregulation is 1 and downregulation is -1.

**Figure 6.**
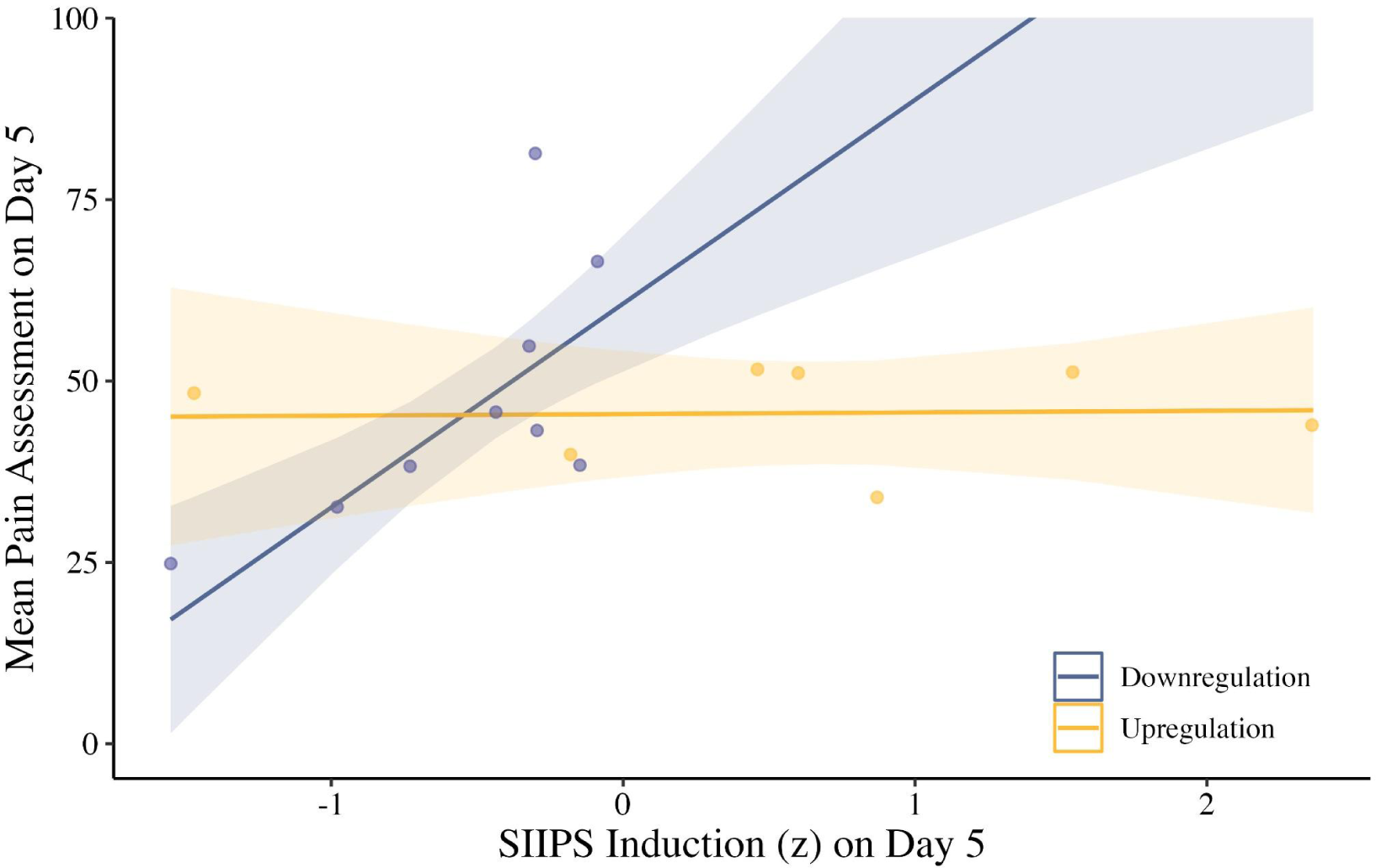
Mean pain assessment predictions on day 5 across SIIPS induction for the upregulation (yellow) and downregulation (blue) groups. Data points indicate the mean pain ratings and SIIPS induction for each individual participant. SIIPS induction values are *z*-scored. Shaded regions indicate standard error.

### The effect of expectations

While our experiment was designed to be conducted in a double-blind fashion (i.e., experimenters and participants were not aware of group membership at the time of the experiment), participants still generally believed they were in the downregulation group when asked at the end of the experiment. More precisely, the false-alarm rate in a two-alternative forced-choice question was found to be .86 for the upregulation group and .11 for the downregulation group. However, this is unlikely to explain the results as both groups equally believed they were attempting to downregulate pain-related brain activity (i.e., 8/9 for the downregulation and 6/7 for the upregulation group).

## Discussion

In this study, we examined whether the subjective experience of pain could be changed by training participants to modulate their own pain-related brain activity (i.e., SIIPS expression). First, we found that pain ratings can be predicted in real-time using a novel method for online decoding in the MNI space. Second, we demonstrated that participants can be trained using decoded neurofeedback to downregulate the expression of SIIPS, which can be achieved independently from NPS expression. Third, we found that the success of SIIPS downregulation was associated with pain perception on Day 5. Taken together, these results suggest that modulating SIIPS expression can in turn change the subjective experience of pain, potentially making it a suitable target for therapeutic interventions.

In order to use an “off-the-shelf” brain decoder, we created a real-time analytical pipeline leveraging GPU computing. Using this approach, decoding in the MNI space could be achieved in approximately .78 seconds on a relatively standard desktop computer. While real-time decoding of whole-brain activity is not new in the native space [23,24], facilitating real-time decoding in the MNI-space might potentially open up a range of new interesting applications. For instance, multiple “off-the-shelf” decoders have previously been trained to predict subjective and physiological outcomes including fear [25,26], negative emotions [16], guilt [27], empathy [28], threat conditioning [29], skin conductance reactivity and autonomic responses [25,30]. Conducting MNI space neurofeedback could potentially help validate such decoders by directly studying the link between the decoders’ expression and their predicted outcome. If this association can be demonstrated in double-blind experiments, it can represent a strong scientific demonstration of the role of the targeted representation in generating the studied outcome.

Our results also suggest that the expression of SIIPS seems to be controlled independently from the expression of NPS. Since both decoders are thought to reflect independent but related processes in pain perception, these results highlight the high level of specificity that can be achieved by decoded neurofeedback interventions [5,12]. Previously, it was suggested that this specificity might be achieved by a mechanism of reinforcement learning occurring at the level of principal components in the brain [9]. In this view, providing a reward that is contingent on the activity of specific principal components might lead, over time, to the specific activation of the targeted brain activity. However, some questions remain regarding the precise mechanisms of action involved in decoded neurofeedback. For instance, it is still unclear why the capacity to down- or up-regulate SIIPS should vary. One possibility is that our experimental design might simply have lacked power in order to reveal the up-regulation of SIIPS during training. Further studies conducted with a greater number of participants will be required in order to answer this question.

We chose to target SIIPS rather than NPS because NPS primarily reflects the intensity of the nociceptive input and might be harder to modulate using decoded neurofeedback [1]. However, this question could be directly investigated in future studies. One possibility is that training participants to modulate NPS might indeed influence the processing of stimulus intensity in the brain and, in turn, affect pain experience. Yet another possibility is that modulating NPS expression may have no direct influence over the subjective experience. Our previous work with fear suggests that the latter scenario is a likely possibility [6,17,31] and future research will be needed in order to fully assess this possibility.

MNI-space decoders present the advantage of avoiding training a different brain decoder for each participant, which can be both lengthy and expensive. However, training accurate MNI-space decoders may not always be possible. For instance, previous work indicates that MNI space decoders of visual processes in the ventral visual stream tend to present relatively low accuracies [14]. This suggests that all patterns of brain activity may not all be a good target for MNI space decoding. The approach might be limited to patterns of brain activity presenting low between-subject variability once they are smoothed and transformed to the MNI space. But, as our results indicate, when this condition is met MNI space decoders might afford similar levels of efficacy as within-subject decoders for decoded neurofeedback purposes. As such, these decoders might represent a good alternative avenue to explore in order to facilitate the implementation of future decoded neurofeedback studies.

In this study, we had to provide feedback for the output of a brain decoder that is not naturally bound. In hopes of providing meaningful feedback to participants, we computed how much the current SIIPS activity deviated from a normative sample estimated from the resting state data of participants. This novel scaling approach was used in order to address the difficulties of interpreting the SIIPS decoder output. As we show, this approach was successful in providing control over SIIPS (Figure 4). Future studies could explore means of improving over this approach. For instance, one might consider instead providing feedback based on the distribution of decoder expression during painful stimulations.

The success of our decoded fMRI neurofeedback intervention may have important implications in designing alternative treatment strategies for patients suffering from pain disorders. Although pain is one of the primary reasons why individuals seek health care [32], a significant portion of individuals with pain-related conditions are not receiving relief from current treatments [33]. Decoded neurofeedback presents a promising alternative because it may be able to change the representation of pain directly and unconsciously in the human brain. As demonstrated by the current study, decoded neurofeedback modulated both the expression of SIIPS and the subjective pain ratings. Even if pain is controlled by current treatments, the undesired side effects may prolong recovery and severely impact the patient’s quality of life [34]. Decoded neurofeedback operates unconsciously, therefore bypassing potential problems with treatment expectations and common side-effects of pain treatments. Taken together, integrating decoded neurofeedback into current interventions may aid in improving the efficacy of treatments for individuals with pain disorders.

Despite these strengths, some limitations should be noted. The sample size in the present study was small, with only 16 participants tested. Small sample sizes can reduce both the statistical power of the study and the reproducibility of results [35]. Therefore, future research would benefit from studies that include larger pre-registered samples to determine if the effect on pain found in our current study can be replicated. A larger sample size would also allow to have more generalizable multilevel models, which may include more random effects. Additionally, although we demonstrated that decoded neurofeedback could achieve some control over the expression of SIIPS, our procedure is still suboptimal. For instance, we were expecting the effect of decoded neurofeedback to be specific to the situation where dot motion also appears on the screen. Instead, we observed a non-specific effect over pain ratings (No main effect or interaction with Condition in Table 3) that could be more closely related to a general effect of priming than to associative learning *per se*. Future studies could study this possibility by investigating the duration of the effect of neurofeedback on pain ratings. Furthermore, future studies could explore new ways of achieving better control over the targeted brain representation [36,37]. This could notably be achieved by exploring new means of scaling and providing feedback to participants.

One puzzling finding is the absence of effect observed in the up-regulation group. One possibility is that there is actually an effect that we simply could not reveal with our design. This is a likely possibility since our results indicate a trend towards significance. The expectation of a down-regulation effect in the up-regulation group may have also counteracted the effect of neurofeedback, making it harder to reveal a statistically significant effect. Yet, another possibility is that there exists a real difference in the capacity to up- or down-regulate SIIPS and that participants may not learn to up-regulate SIIPS reliably. Since asymmetrical results have sometimes been observed in other neurofeedback studies [31], a replication study might be needed to address this question by including a broader up-regulation group and an experimental manipulation of the expectations of participants.

Overall, this study demonstrates, in a rigorous experimental design, that decoded fMRI neurofeedback can modulate the subjective experience of pain. These findings yield valuable knowledge to aid in designing future interventions for patients suffering from pain disorders. Currently, there is limited research examining the power of real-time brain imaging in manipulating affective experiences [6,7]. Future studies should aim to replicate and extend the current findings to better understand how control over affective processes can be achieved. Our novel additions to the decoded neurofeedback methodology should allow future studies to expand and build upon this literature by allowing one to easily target MNI-space decoders. These innovations could help actualise the promises of decoded neurofeedback to be used more readily in clinical settings as a complementary psychological intervention.

## Acknowledgements, funding statement and disclosure

This research was conducted with the financial support of the Tom Slick research award in consciousness 2020 from the Mind Science foundation. V.T-D. was supported in part by the Canadian Institute of Health Research and the *Fond de recherche du Québec - Santé*. A.C. and M.K. are partially supported by the Japan Science and Technology agency ERATO Ikegaya brain-AI fusion (grant number JPJMER1801), by JSPS KAKENHI (grant number JP22H05156), and by the Agency for Technology, Labour and Innovation (grant number JP004596). Author Mitsuo Kawato is an inventor of patents owned by the Advanced Telecommunications Research Institute International related to the present work (PCT/JP2012/078136 [WO2013/06 871 9517] and PCT/JP2014/61543 [WO2014/178322]). The other authors have no conflicts to declare.

## Supplementary Materials

**Table S1.** *Multilevel model estimates for decoder induction. Estimates are bootstrapped 10 000 times.*

| *Predictors* | *Model Estimates* | *95% CI* | *p-values* | *Bootstrapped Estimates* | *Bootstrapped 95% CI* | *p-values* |
| --- | --- | --- | --- | --- | --- | --- |
| (Intercept) | 537.807 | [-17.334, 1092.948] | .058 | 539.855 | [-14.675, 1107.700] | .056 |
| Trial | 28.608 | [12.475, 44.740] | <.001 | 28.656 | [12.475, 44.842] | <.001 |
| log(Trial) | -859.382 | [-1474.588, -244.176] | .006 | -860.561 | [-1469.599, -244.421] | .006 |
| Group | 165.949 | [-384.909, 716.807] | .555 | 168.237 | [-373.337, 710.468] | .555 |
| Day | 61.066 | [-238.584, 360.716] | .690 | 61.327 | [-241.867, 364.559] | .684 |
| Decoder Type | 575.968 | [137.345, 1014.591] | .010 | 580.615 | [149.498, 1014.651] | .012 |
| Group x Day | 349.974 | [56.355, 643.593] | .019 | 353.599 | [58.433, 640.564] | .019 |
| Group x Decoder Type | 155.017 | [-283.606, 593.640] | .488 | 151.731 | [-287.244, 589.515] | .495 |
| Day x Decoder Type | 26.787 | [-266.798, 320.371] | .858 | 27.368 | [-259.170, 317.728] | .859 |
| Group x Day x Decoder Type | 355.742 | [62.158, 649.326] | .018 | 358.141 | [66.942, 650.002] | .016 |
| *Random Effects* |  |  |  |  |  |  |
| $\sigma^2$ | 262792755.53 | | | | | |
| $\tau_0^2$ | 454993.80 | | | | | |
| Marginal/<br>Conditional $R^2$ | .005/ .007 | | | | | |

*Note.* Group is effect coded where upregulation is 1 and downregulation is -1. Decoder type: SIIPS (1) and NPS (-1)

**Table S2.**

| *Predictors* | *Model Estimates* | *95% CI* | *p-values* | *Bootstrapped Estimates* | *Bootstrapped 95% CI* | *p-values* |
| --- | --- | --- | --- | --- | --- | --- |
| (Intercept) | 808.544 | [-239.565, 1856.654] | .164 | 807.525 | [-222.701, 1860.547] | .128 |
| Trial | 14.676 | [1.683, 27.668] | .027 | 14.744 | [1.631, 27.738] | .028 |
| log(Trial) | -391.220 | [-887.949, 105.508] | .123 | -391.449 | [-898.474, 112.763] | .126 |
| Day | -630.537 | [-871.207, -389.867] | <.001 | -632.032 | [-868.812, -385.656] | <.001 |
| *Random Effects* |  |  |  |  |  |  |
| $\sigma^2$ | 48385848.15 | | | | | |
| $\tau_0^2$ | 2276303.85 | | | | | |
| Marginal/<br>Conditional $R^2$ | .009/ .054 | | | | | |

**Table S3.**

| *Predictors* | *Model Estimates* | *95% CI* | *p-values* | *Bootstrapped Estimates* | *Bootstrapped 95% CI* | *p-values* |
| --- | --- | --- | --- | --- | --- | --- |
| (Intercept) | 1354.016 | [-780.004, 3488.036] | .225 | 1363.166 | [-769.601, 3463.894] | .211 |
| Trial | 113.210 | [40.933, 185.487] | .002 | 114.152 | [42.233, 186.722] | .001 |
| log(Trial) | -3461.067 | [-6208.277, -713.857] | .014 | -3263.528 | [-6263.641, -673.236] | .012 |
| Day | 871.926 | [-451.828, 2195.690] | .197 | 875.656 | [-442.724, 2194.109] | .199 |
| *Random Effects* |  |  |  |  |  |  |
| $\sigma^2$ | 1138615295.96 | | | | | |
| $\tau_0^2$ | 1249368.08 | | | | | |
| Marginal/<br>Conditional<br>R<sup>2</sup> | .004/ .005 |  |  |  |  |  |

**Table S4.**
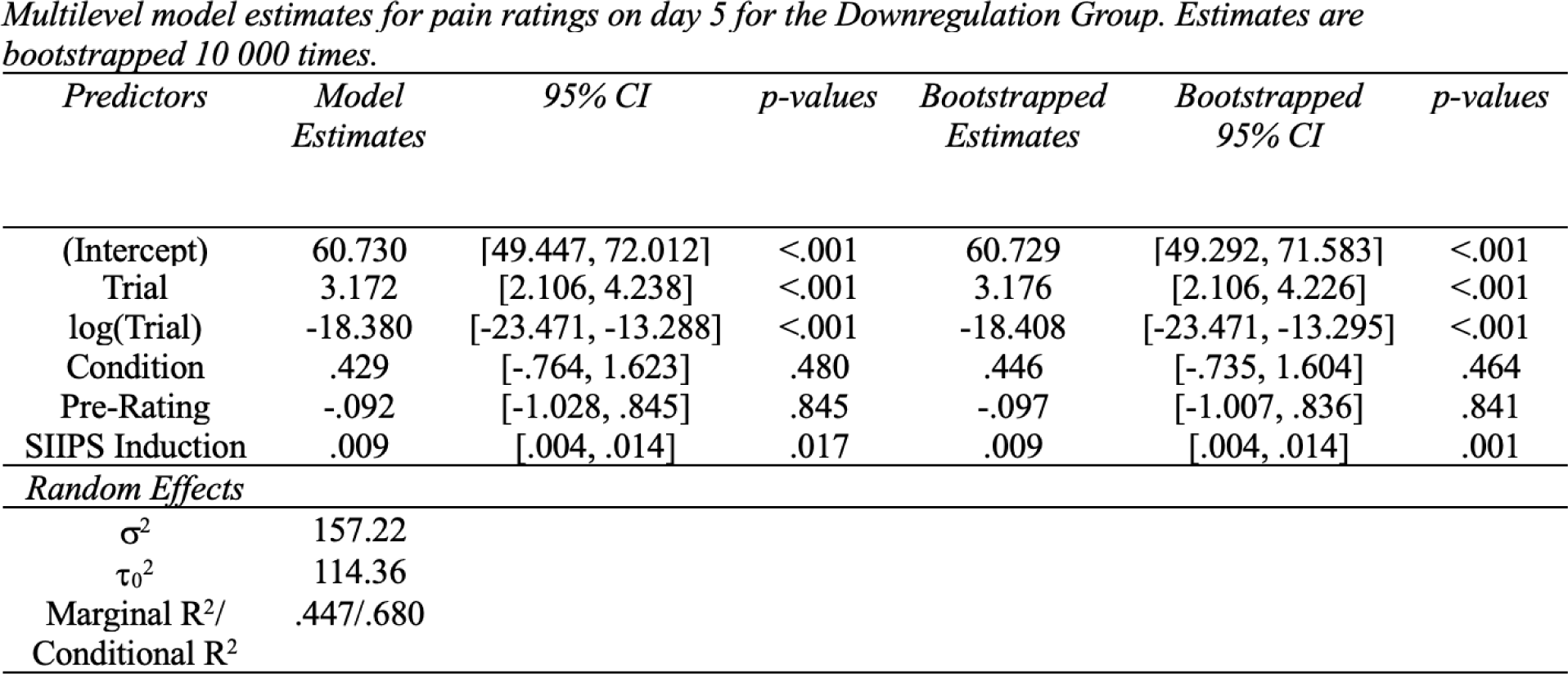

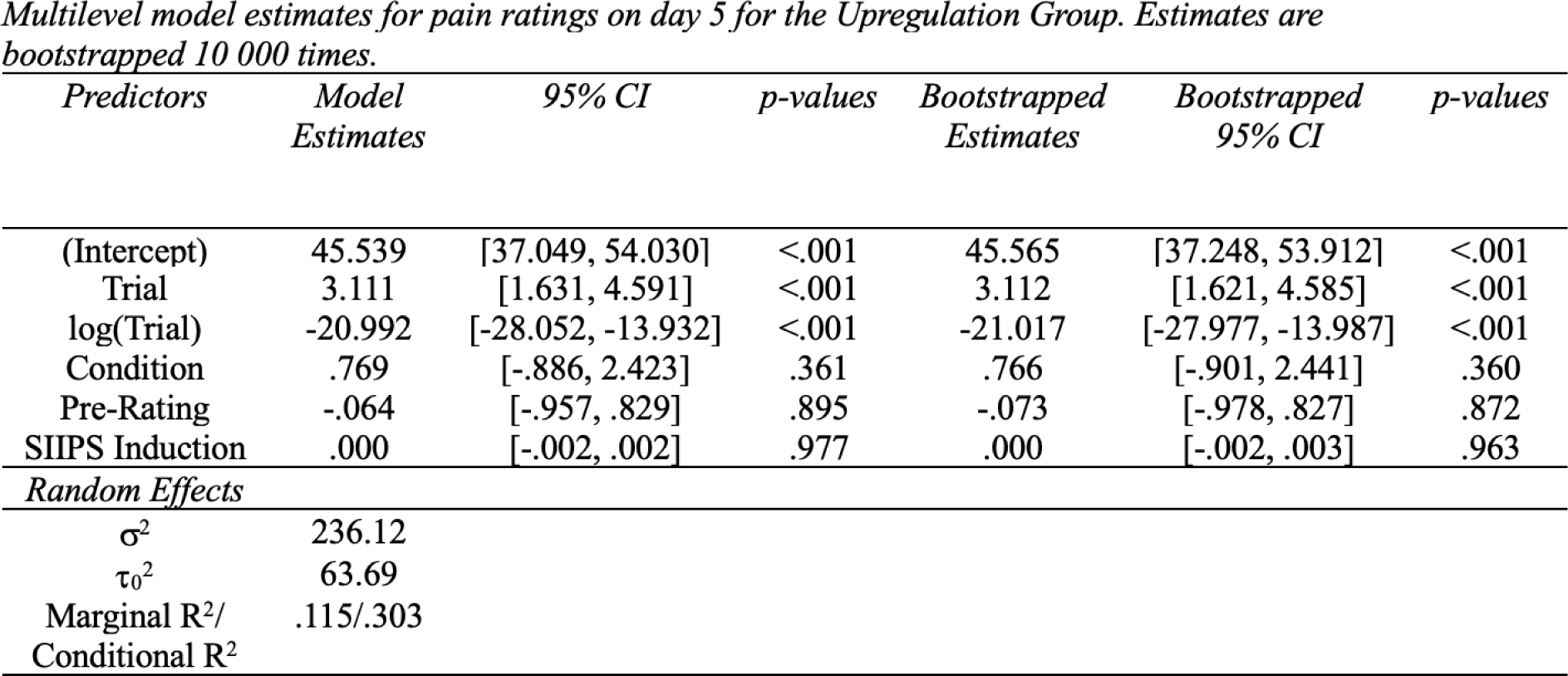

| *Predictors* | *Model Estimates* | *95% CI* | *p-values* | *Bootstrapped Estimates* | *Bootstrapped 95% CI* | *p-values* |
| --- | --- | --- | --- | --- | --- | --- |
| (Intercept) | 60.730 | [49.447, 72.012] | <.001 | 60.729 | [49.292, 71.583] | <.001 |
| Trial | 3.172 | [2.106, 4.238] | <.001 | 3.176 | [2.106, 4.226] | <.001 |
| log(Trial) | -18.380 | [-23.471, -13.288] | <.001 | -18.408 | [-23.471, -13.295] | <.001 |
| Condition | .429 | [-.764, 1.623] | .480 | .446 | [-.735, 1.604] | .464 |
| Pre-Rating | -.092 | [-1.028, .845] | .845 | -.097 | [-1.007, .836] | .841 |
| SIIPS Induction | .009 | [.004, .014] | .017 | .009 | [.004, .014] | .001 |
| *Random Effects* |  |  |  |  |  |  |
| $\sigma^2$ | 157.22 | | | | | |
| $\tau_0^2$ | 114.36 | | | | | |
| Marginal R<sup>2</sup>/<br>Conditional R<sup>2</sup> | .447/.680 |  |  |  |  |  |
| (Intercept) | 45.539 | [37.049, 54.030] | <.001 | 45.565 | [37.248, 53.912] | <.001 |
| Trial | 3.111 | [1.631, 4.591] | <.001 | 3.112 | [1.621, 4.585] | <.001 |
| log(Trial) | -20.992 | [-28.052, -13.932] | <.001 | -21.017 | [-27.977, -13.987] | <.001 |
| Condition | .769 | [-.886, 2.423] | .361 | .766 | [-.901, 2.441] | .360 |
| Pre-Rating | -.064 | [-.957, .829] | .895 | -.073 | [-.978, .827] | .872 |
| SIIPS Induction | .000 | [-.002, .002] | .977 | .000 | [-.002, .003] | .963 |
| *Random Effects* |  |  |  |  |  |  |
| $\sigma^2$ | 236.12 | | | | | |
| $\tau_0^2$ | 63.69 | | | | | |
| Marginal R<sup>2</sup>/<br>Conditional R<sup>2</sup> | .115/.303 |  |  |  |  |  |

## References

1. Woo C-W, Schmidt L, Krishnan A, Jepma M, Roy M, Lindquist MA, Atlas LY, Wager TD. 2017 Quantifying cerebral contributions to pain beyond nociception. Nat. Commun. 8, 14211.

2. Wager TD, Atlas LY, Lindquist MA, Roy M. 2013 An fMRI-based neurologic signature of physical pain. J. Med.

3. Watanabe T, Sasaki Y, Shibata K, Kawato M. 2017 Advances in fMRI Real-Time Neurofeedback. Trends Cogn. Sci. 21, 997–1010.

4. Amano K, Shibata K, Kawato M, Sasaki Y, Watanabe T. 2016 Learning to Associate Orientation with Color in Early Visual Areas by Associative Decoded fMRI Neurofeedback. Curr. Biol. 26, 1861–1866.

5. Shibata K, Watanabe T, Sasaki Y, Kawato M. 2011 Perceptual learning incepted by decoded fMRI neurofeedback without stimulus presentation. Science 334, 1413–1415.

6. Taschereau-Dumouchel V, Cortese A, Chiba T, Knotts JD, Kawato M, Lau H. 2018 Towards an unconscious neural reinforcement intervention for common fears. Proc. Natl. Acad. Sci. U. S. A. 115, 3470–3475.

7. Koizumi A, Amano K, Cortese A, Shibata K, Yoshida W, Seymour B, Kawato M, Lau H. 2016 Fear reduction without fear through reinforcement of neural activity that bypasses conscious exposure. Nat Hum Behav 1. (doi:10.1038/s41562-016-0006)

8. Cortese A, Amano K, Koizumi A, Kawato M, Lau H. 2016 Multivoxel neurofeedback selectively modulates confidence without changing perceptual performance. Nat. Commun. 7, 13669.

9. Shibata K, Lisi G, Cortese A, Watanabe T, Sasaki Y, Kawato M. 2019 Toward a comprehensive understanding of the neural mechanisms of decoded neurofeedback. Neuroimage 188, 539–556.

10. Taschereau-Dumouchel V, Liu K-Y, Lau H. 2018 Unconscious Psychological Treatments for Physiological Survival Circuits. Curr Opin Behav Sci 24, 62–68.

11. Taschereau-Dumouchel V, Cortese A, Lau H, Kawato M. 2020 Conducting Decoded Neurofeedback Studies. Soc. Cogn. Affect. Neurosci. (doi:10.1093/scan/nsaa063)

12. Shibata K, Watanabe T, Kawato M, Sasaki Y. 2016 Differential Activation Patterns in the Same Brain Region Led to Opposite Emotional States. PLoS Biol. 14, e1002546.

13. deCharms RC, Maeda F, Glover GH, Ludlow D, Pauly JM, Soneji D, Gabrieli JDE, Mackey SC. 2005 Control over brain activation and pain learned by using real-time functional MRI. Proceedings of the National Academy of Sciences 102, 18626–18631.

14. Haxby JV, Guntupalli JS, Connolly AC, Halchenko YO, Conroy BR, Gobbini MI, Hanke M, Ramadge PJ. 2011 A common, high-dimensional model of the representational space in human ventral temporal cortex. Neuron 72, 404–416.

15. Lindquist MA et al. 2017 Group-regularized individual prediction: theory and application to pain. Neuroimage 145, 274–287.

16. Chang LJ, Gianaros PJ, Manuck SB, Krishnan A, Wager TD. 2015 A Sensitive and Specific Neural Signature for Picture-Induced Negative Affect. PLoS Biol. 13, e1002180.

17. Taschereau-Dumouchel V, Michel M, Lau H, Hofmann SG, LeDoux JE. 2022 Putting the ‘mental’ back in ‘mental disorders’: a perspective from research on fear and anxiety. Mol. Psychiatry (doi:10.1038/s41380-021-01395-5)

18. Eklund A, Dufort P, Villani M, Laconte S. 2014 BROCCOLI: Software for fast fMRI analysis on many-core CPUs and GPUs. Front. Neuroinform. 8, 24.

19. Penny WD, Friston KJ, Ashburner JT, Kiebel SJ, Nichols TE. 2011 Statistical Parametric Mapping: The Analysis of Functional Brain Images. Elsevier.

20. Lüdecke D. 2021 sjPlot: Data Visualization for Statistics in Social Science.(Version 2.8. 10)[Computer software].

21. Team R. 2020 RStudio: integrated development for R. RStudio, PBC, Boston, MA.

22. Bates D, Mächler M, Bolker B, Walker S. 2014 Fitting Linear Mixed-Effects Models using lme4. arXiv,1406.5823.

23. deBettencourt MT, Cohen JD, Lee RF, Norman KA, Turk-Browne NB. 2015 Closed-loop training of attention with real-time brain imaging. Nat. Neurosci. 18, 470–475.

24. Mennen AC et al. 2021 Cloud-Based Functional Magnetic Resonance Imaging Neurofeedback to Reduce the Negative Attentional Bias in Depression: A Proof-of-Concept Study. Biol Psychiatry Cogn Neurosci Neuroimaging 6, 490–497.

25. Taschereau-Dumouchel V, Kawato M, Lau H. 2020 Multivoxel pattern analysis reveals dissociations between subjective fear and its physiological correlates. Mol. Psychiatry 25, 2342–2354.

26. Zhou F, Zhao W, Qi Z, Geng Y, Yao S. 2021 A distributed fMRI-based signature for the subjective experience of fear. Nature Communications

27. Yu H, Koban L, Crockett MJ, Zhou X, Wager TD. 2020 Toward a Brain-Based Bio-Marker of Guilt. Neurosci Insights. 15, 2633105520957638.

28. Ashar YK, Andrews-Hanna JR, Dimidjian S, Wager TD. 2017 Empathic Care and Distress: Predictive Brain Markers and Dissociable Brain Systems. Neuron 94, 1263–1273.e4.

29. Reddan MC, Wager TD, Schiller D. 2018 Attenuating Neural Threat Expression with Imagination. Neuron 100, 994–1005.e4.

30. Eisenbarth H, Chang LJ, Wager TD. 2016 Multivariate Brain Prediction of Heart Rate and Skin Conductance Responses to Social Threat. J. Neurosci. 36, 11987–11998.

31. Cushing CA, Lau H, Kawato M, Craske MG, Taschereau-Dumouchel V. 2023 A pre-registered decoded neurofeedback intervention for specific phobias. medRxiv, 2023–2004.

32. Finley CR et al. 2018 What are the most common conditions in primary care?: Systematic review. Can. Fam. Physician 64, 832–840.

33. Sinatra R. 2010 Causes and consequences of inadequate management of acute pain. Pain Med. 11, 1859–1871.

34. Chabot B, Ferland CE. 2020 Inpatient postoperative undesirable side effects of analgesics management: a pediatric patients and parental perspective. Pain Rep 5, e845.

35. Button KS, Ioannidis JPA, Mokrysz C, Nosek BA, Flint J, Robinson ESJ, Munafò MR. 2013 Power failure: why small sample size undermines the reliability of neuroscience. Nat. Rev. Neurosci. 14, 365–376.

36. Taschereau-Dumouchel V, Cushing C, Lau H. 2022 Real-time functional MRI in the treatment of mental health disorders. Annual review of clinical psychology

37. Taschereau-Dumouchel V, Roy M. 2020 Could Brain Decoding Machines Change Our Minds? Trends Cogn. Sci. (doi:10.1016/j.tics.2020.09.006)

